## Supplementary Information for "On the discovery of subpopulation-specific state transitions from multi-sample multi-condition single-cell RNA sequencing data"

| Method ID | Input data | Agg. | Model level | Ref. |
|---|---|---|---|---|
| edgeR.sum(counts) | counts | ✓ | cluster-sample PBs | Robinson *et al.*[1] |
| edgeR.sum(scalecpm) | LS scaled pseudobulk CPM | ✓ | cluster-sample PBs | |
| limma-voom.sum(counts) | counts | ✓ | cluster-sample PBs | Ritchie *et al.*[2] |
| limma-trend.mean(logcounts) | $\log_2$ LS normalized counts | ✓ | cluster-sample PBs | |
| limma-trend.mean(vstresiduals) | VST residuals | ✓ | cluster-sample PBs | |
| MM-dream | counts | ✗ | SCs; cluster-level | Hoffman & Schadt[3] |
| MM-dream2 | counts | ✗ | SCs; cluster-level | **Hoffman2020-variancePartition**[Hoffman2020-variancePartition] |
| MM-nbinom | counts | ✗ | SCs; cluster-level | |
| MM-vst | VST residuals | ✗ | SCs; cluster-level | |
| scDD.logcounts | $\log_2$ LS normalized counts | ✗ | SCs; cluster-level | Korthauer *et al.*[4] |
| scDD.vstresiduals | VST residuals | ✗ | SCs; cluster-level | |
| MAST.logcounts | $\log_2$ LS normalized counts | ✗ | SCs; cluster-level | Finak *et al.*[5] |
| AD-gid.logcounts | $\log_2$ LS normalized counts | ✗ | SCs; cluster-group level | Scholz & Stephens[6] |
| AD-gid.vstresiduals | VST residuals | ✗ | SCs; cluster-group level | |
| AD-sid.logcounts | $\log_2$ LS normalized counts | ✗ | SCs; cluster-sample level | |
| AD-sid.vstresiduals | VST residuals | ✗ | SCs; cluster-sample level | |

**Table 1: Overview of compared DS analysis methods.** From left to right: Method identifier as depicted in all figures; input data; whether data is aggregated or not; the levels at which differential testing is performed; reference. (Agg. = aggregation, CPM = counts per million, LS = library size, VST = variance stabilizing transformation, PBs = pseudobulks, SCs = single cells)

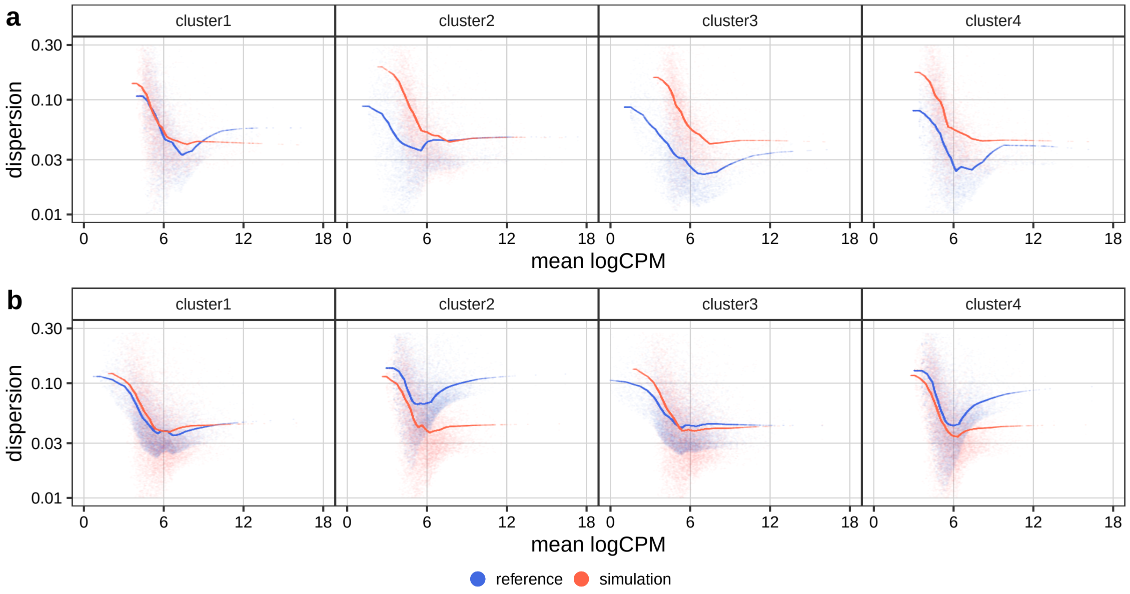


**Supplementary Figure 1: Comparison of pseudobulk-level mean-dispersion estimates for reference vs. simulated data, separated by subpopulation.** Lines correspond to trended dispersion estimates; faded points represent tag-wise dispersion estimates. Lower (1%) and upper (99%) dispersion quantiles were removed for visualization. Simulations are based on the (**a**) Kang *et al.*[7] and (**b**) LPS dataset as reference, respectively.

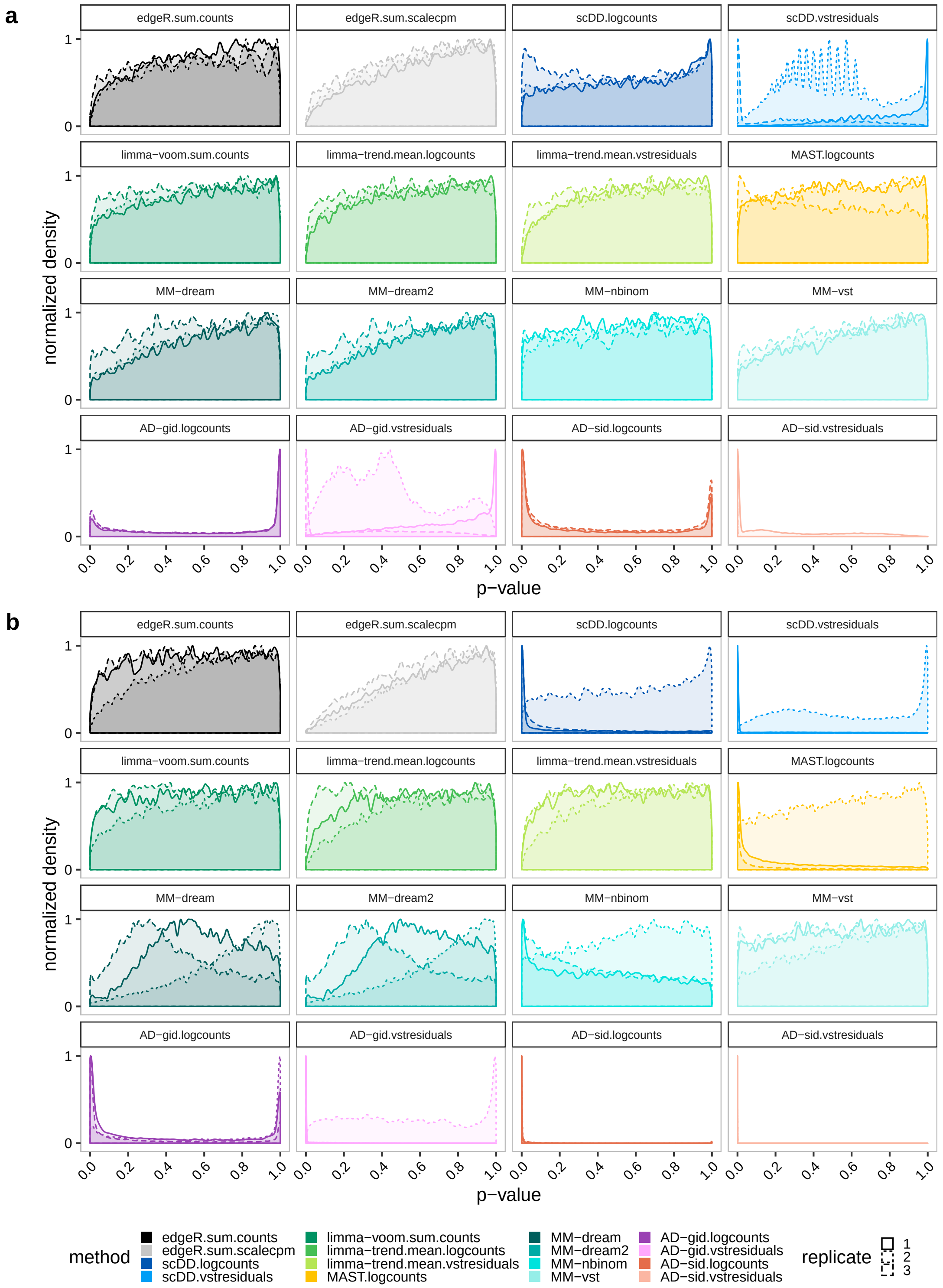


**Supplementary Figure 2: Nominal p-value distributions (densities) obtained from three null simulation replicates, stratified by method.** Each simulation run includes 3 samples per group, and 2000 genes tested across 2 clusters; 3 simulation runs are shown. Densities that are near-uniform are consistent with data lacking differential signal. Simulations are based on the (**a**) Kang *et al.*[7] and (**b**) LPS dataset as reference, respectively.

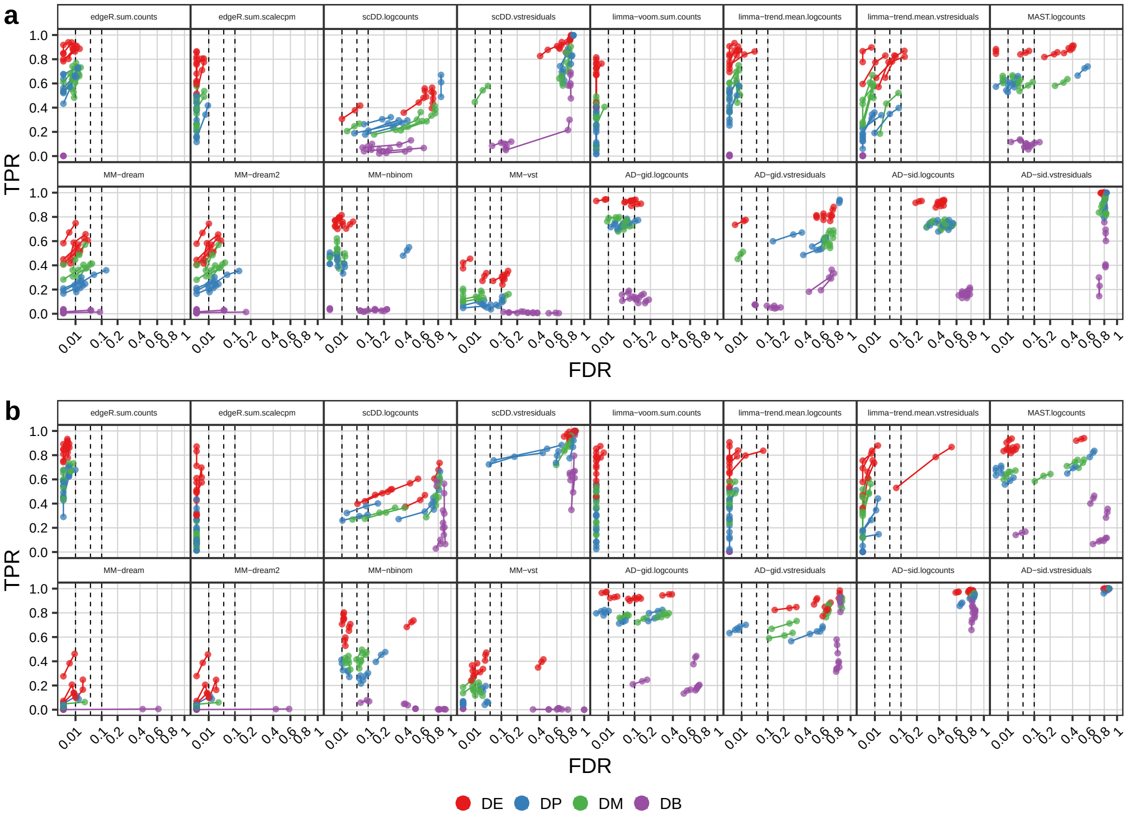


**Supplementary Figure 3: DS method performances across differential distribution and simulation replicates.** Points correspond to observed overall true positive rate (TPR) and false discovery rate (FDR) values at FDR cutoffs of 1%, 5%, and 10%; dashed lines indicate desired FDRs. Each group of inter-connected points corresponds to one simulation with 10% of DS genes (of the type indicated by their color). Simulations are based on the (**a**) Kang *et al.*[7] and (**b**) LPS dataset as reference, respectively.

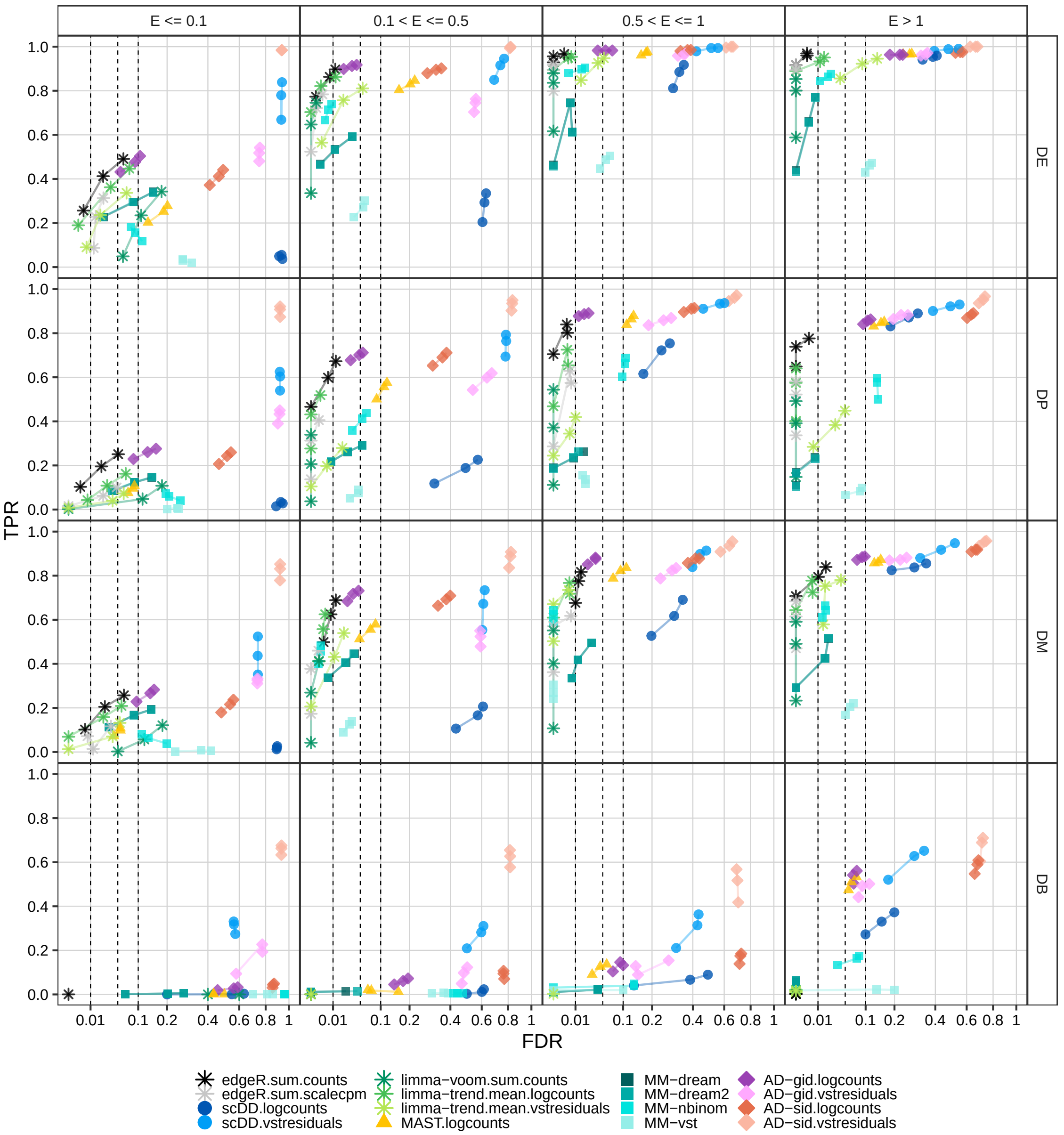


**Supplementary Figure 4: DS method performances across expression-levels and differential distribution categories; Kang et al.[7] dataset reference.** Points correspond to observed overall true positive rate (TPR) and false discovery rate (FDR) values at FDR cutoffs of 1%, 5%, and 10%; dashed lines indicate desired FDRs. Results were stratified into groups according to the mean of simulated expression-means across groups. For each panel, performances were averaged across 5 simulation replicates, each containing 10% of DS genes.

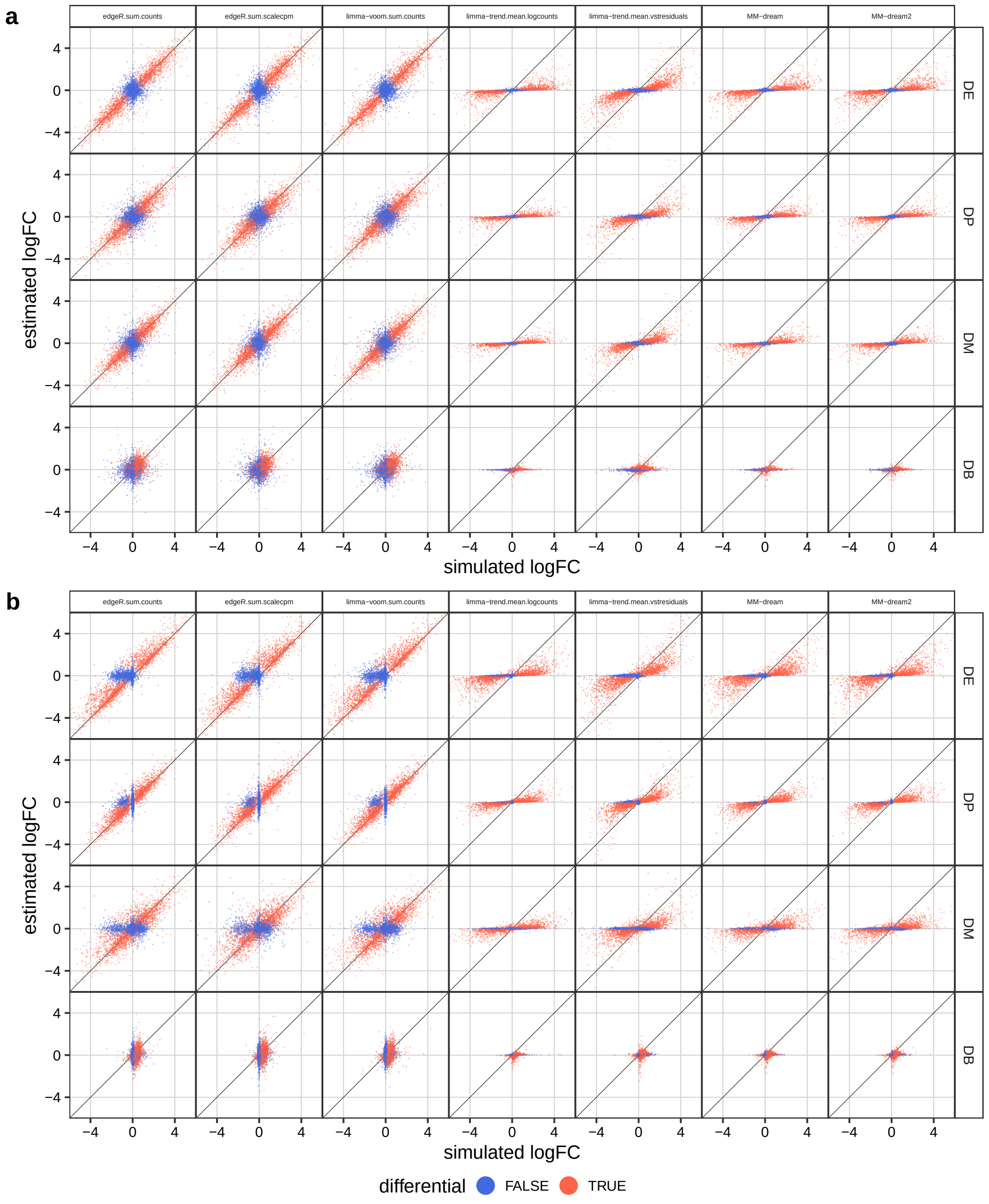


**Supplementary Figure 5: Simulated vs. estimated cross-group log-fold changes (logFC), stratified by method and gene category.** Each point corresponds to a gene-subpopulation instance; coloring corresponds to non-differential (blue) or truly differential (red). Included are only methods that return logFC estimates. For plotting, a random subset of 2'000 points was sampled per method, simulation, and color. Simulations are based on the (**a**) Kang *et al.*[7] and (**b**) LPS dataset as reference, respectively.

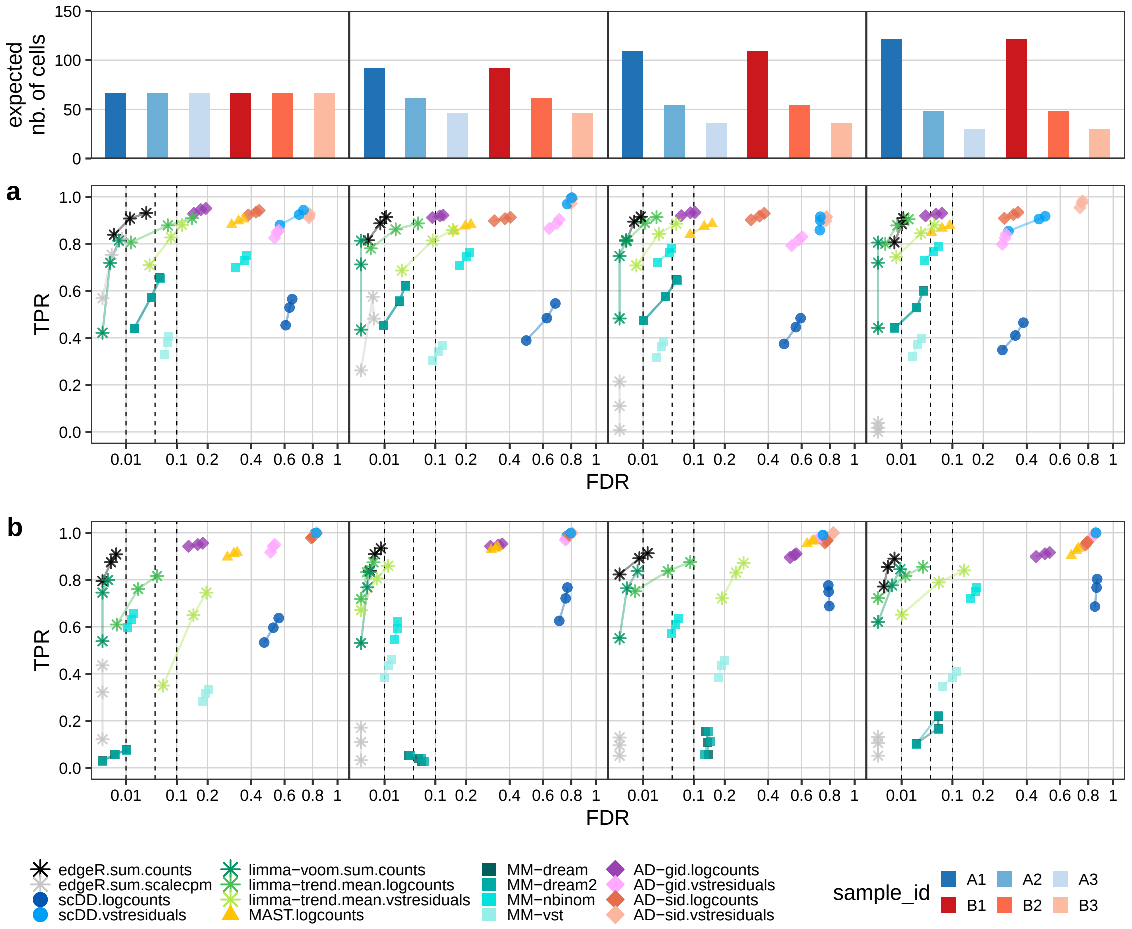


**Supplementary Figure 6: Effects of unbalanced sample sizes on DS method performances.** Points correspond to observed overall true positive rate (TPR) and false discovery rate (FDR) values at FDR cutoffs of 1%, 5%, and 10%; dashed lines indicate desired FDRs. Results were stratified into groups according to the variance of simulated sample sizes. For each panel, performances were averaged across 5 simulation replicates, each containing 10% of DS genes. Simulations are based on the (**a**) Kang *et al.*[7] and (**b**) LPS dataset as reference, respectively.

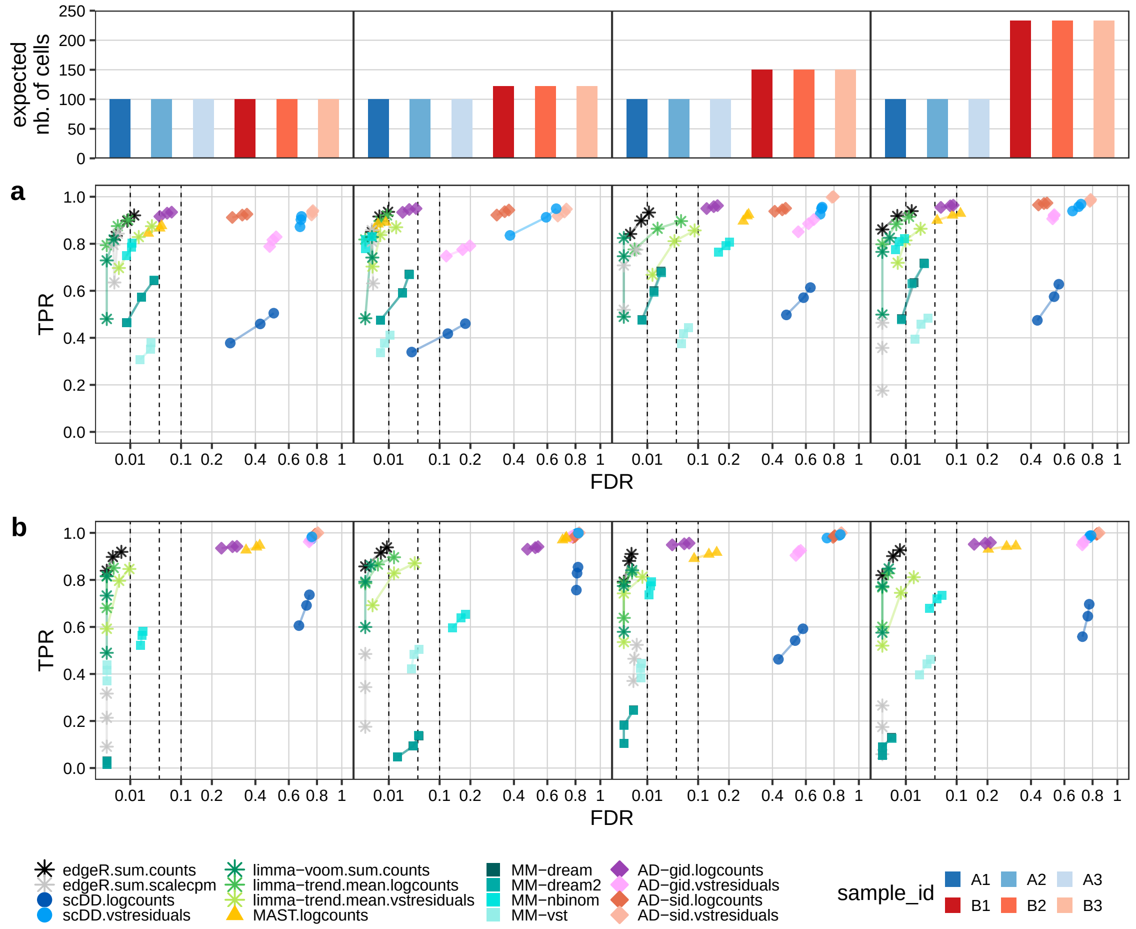


**Supplementary Figure 7: Effects of unbalanced group sizes on DS method performances.** Points correspond to observed overall true positive rate (TPR) and false discovery rate (FDR) values at FDR cutoffs of 1%, 5%, and 10%; dashed lines indicate desired FDRs. Results were stratified into groups according to the variance of simulated group sizes. For each panel, performances were averaged across 5 simulation replicates, each containing 10% of DS genes. Simulations are based on the (**a**) Kang *et al.*[7] and (**b**) LPS dataset as reference, respectively.

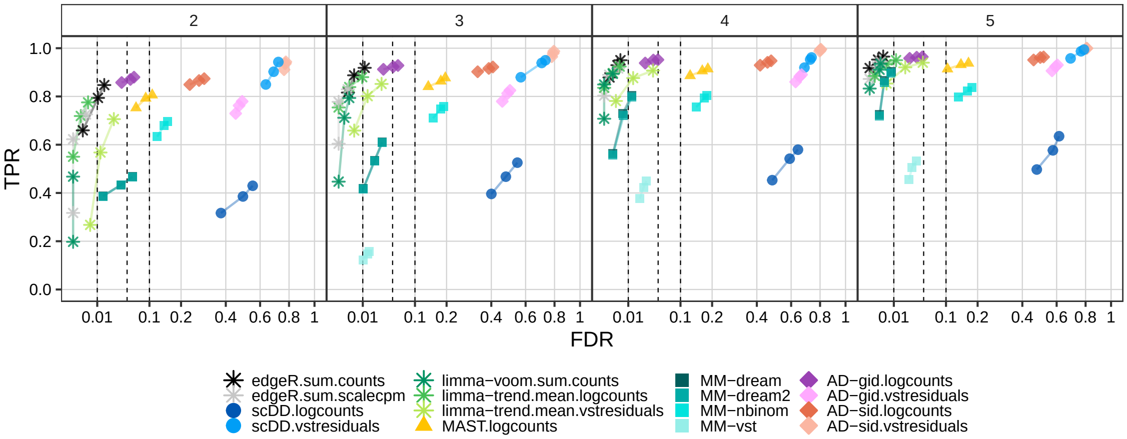


**Supplementary Figure 8: Effect of the number of replicates per group on DS method performances; Kang et al.[7] dataset reference.** Points correspond to observed overall true positive rate (TPR) and false discovery rate (FDR) values at FDR cutoffs of 1%, 5%, and 10%; dashed lines indicate desired FDRs. Results were stratified into groups according to the number replicates in each group. For each panel, performances were averaged across 5 simulation replicates, each containing 10% of DS genes.

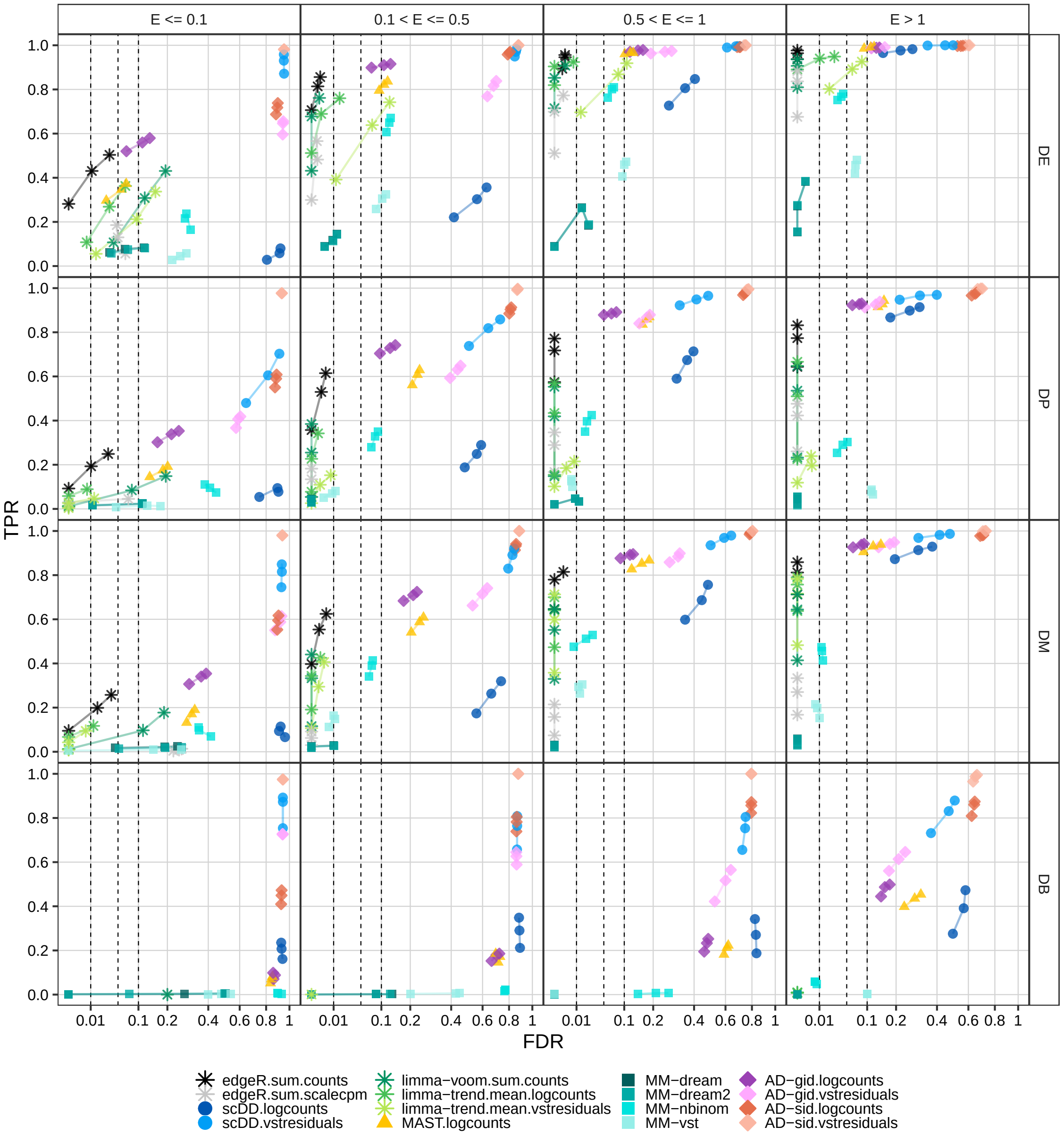


**Supplementary Figure 9: DS method performances across expression levels and differential distribution categories; LPS dataset reference.** Points correspond to observed overall true positive rate (TPR) and false discovery rate (FDR) values at FDR cutoffs of 1%, 5%, and 10%; dashed lines indicate desired FDRs. Results were stratified into groups according to the mean of simulated expression-means across groups. For each panel, performances were averaged across 5 simulation replicates, each containing 10% of DS genes (of the type specified in the right-hand side panel labels).

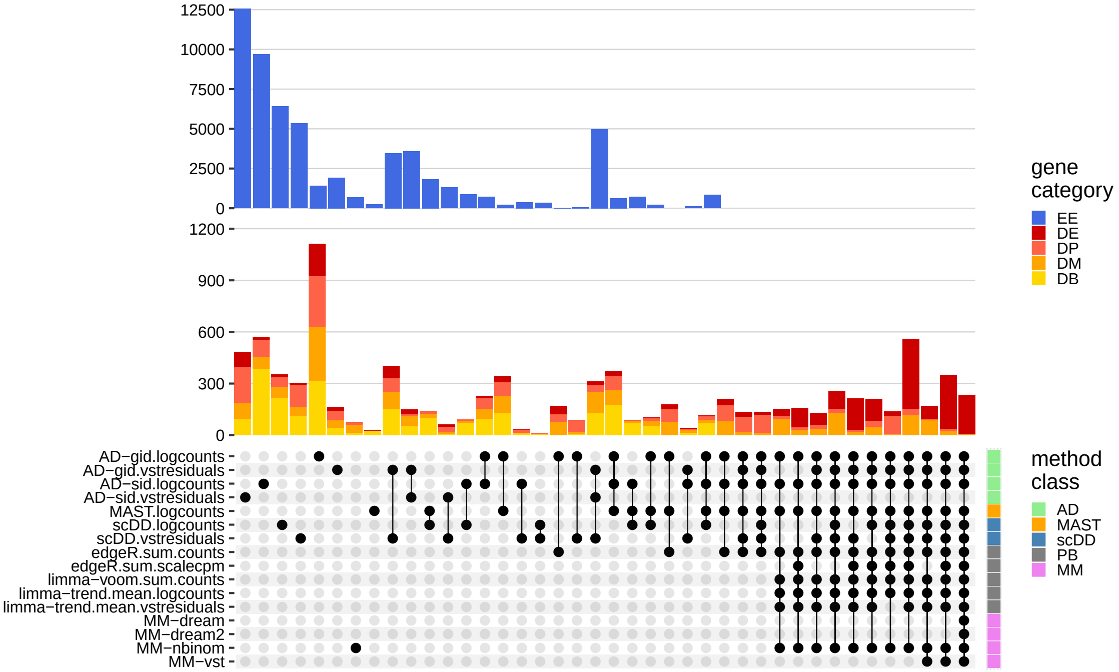


**Supplementary Figure 10: Between-method concordance; LPS dataset reference.** Upset plot obtained from intersecting the top-$n$ ranked differential genes, where $n = min(n_1, n_2)$, where $n_1$ = number of genes simulated to be differential, and $n_2$ = number of genes called differential at FDR $< 0.05$. Shown are the 40 most frequent interactions; coloring corresponds to (true) simulated gene categories.

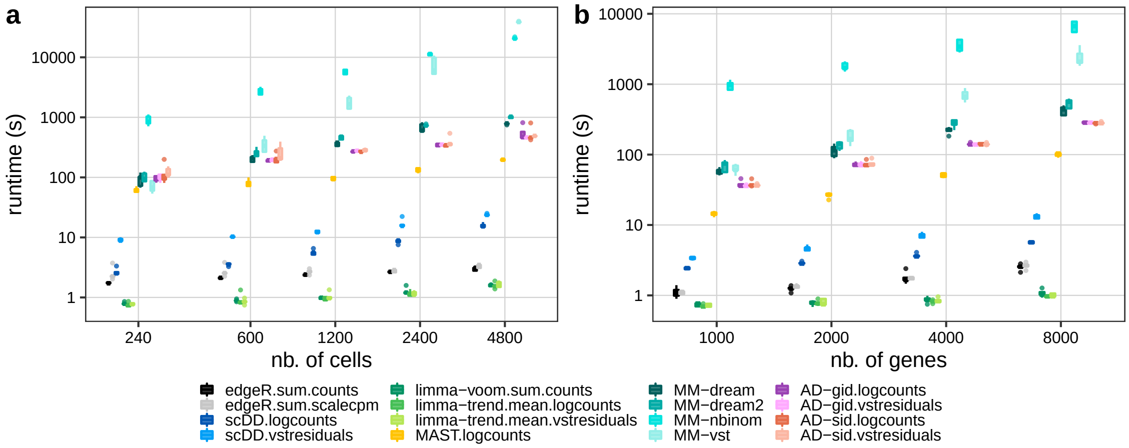


**Supplementary Figure 11: DS method runtimes** vs. (**a**) number of cells and (**b**) number of genes. Included are runtimes from 5 simulation replicates per subset of cells and genes, respectively, using the Kang *et al.*[7] dataset reference; single-core computing times were recorded.

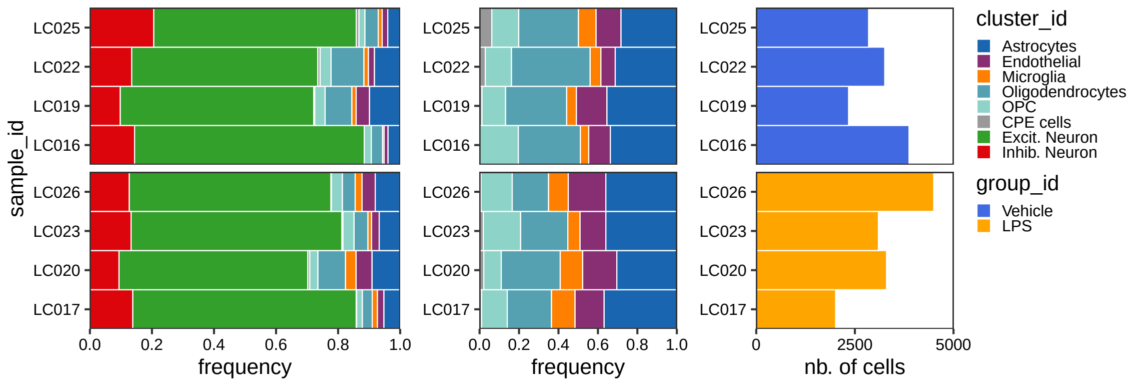


**Supplementary Figure 12: Relative and absolute subpopulation abundances for the LPS dataset.** The left panel shows sample-wise frequencies of the annotated subpopulations; the middle panel shows relative frequencies after removal of all neuronal subpopulations; the right panel shows the number of cells per sample after filtering.

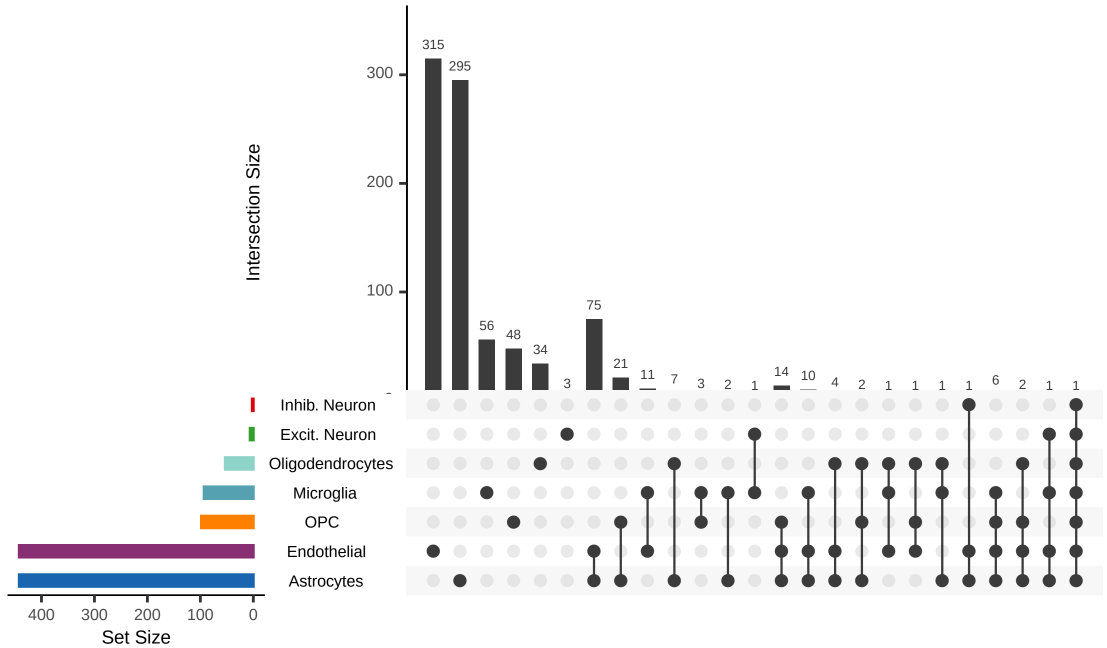


**Supplementary Figure 13: Upset plot of differentially expressed genes identified for the LPS dataset, by detected subpopulation.** Included are genes with FDR $< 0.05$ and $|\text{logFC}| > 1$; shown are all subpopulations intersections with non-zero size.

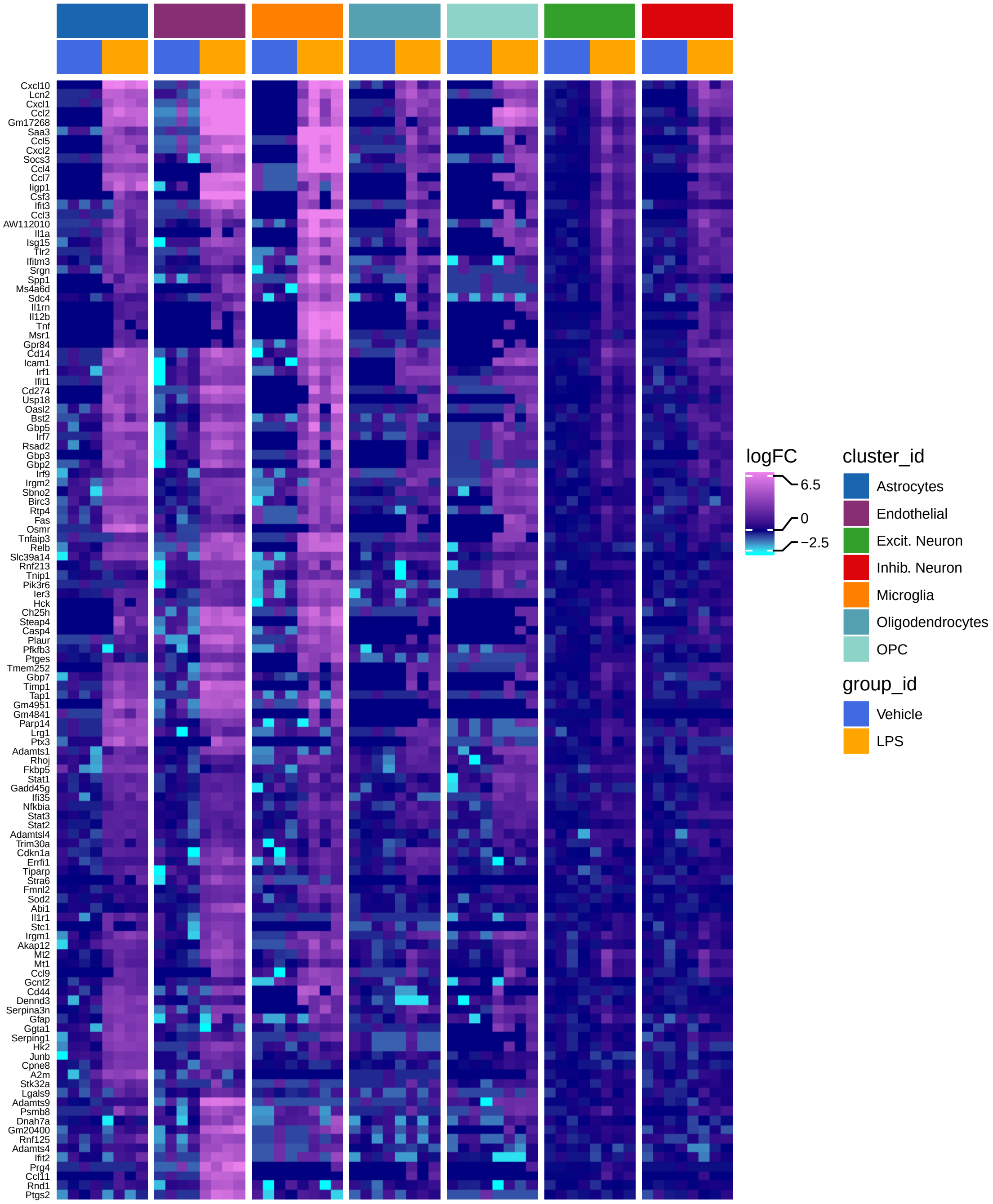


**Supplementary Figure 14: Heatmap of cross-group logFCs of DE genes with consensus cluster ID 3 for the LPS dataset.** Included are DE genes with FDR $< 1\text{e}{-4}$ and $|\text{logFC}| > 1$. For every gene, the displayed log-fold-change (logFC) is normalized to that gene's average expression in the vehicle group (in the corresponding subpopulation); top and bottom 1% logFC quantiles were truncated for visualization.

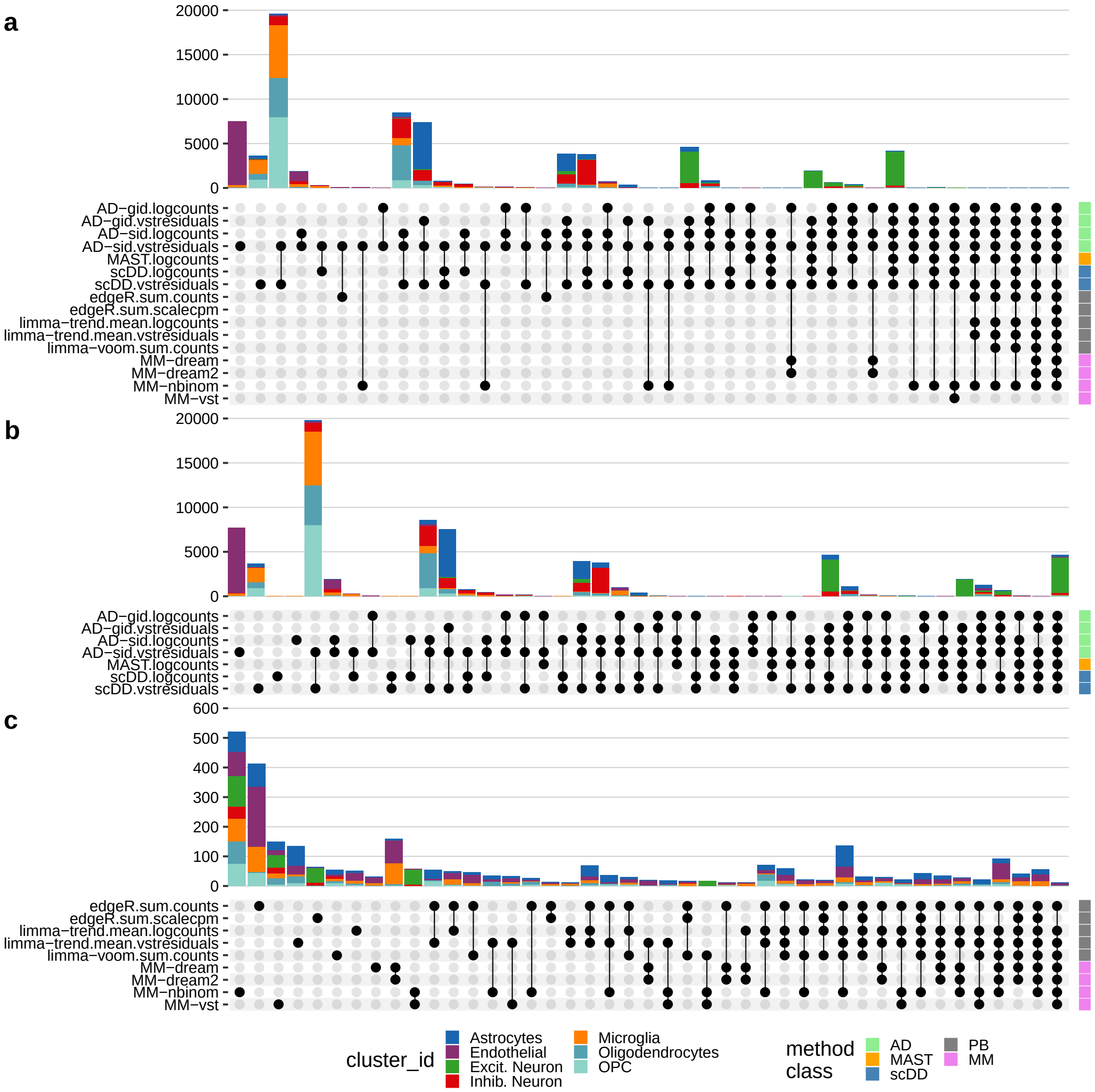


**Supplementary Figure 15: Upset plot of differential state genes detected for the LPS dataset, by method and across all subpopulations (excluding CPE cells).** Included are genes with FDR $< 0.05$; shown are the 40 most frequent intersections between (**a**) all methods, (**b**) AD, MAST and scDD methods, and (**c**) aggregation- and MM-based methods.
